# Optimization of SARS-CoV-2 detection by RT-QPCR without RNA extraction

**DOI:** 10.1101/2020.04.06.028902

**Authors:** Natacha Merindol, Geneviève Pépin, Caroline Marchand, Marylène Rheault, Christine Peterson, André Poirier, Hugo Germain, Alexis Danylo

**Author notes:** contributed equally to this work (Hugo Germain, Alexis Danylo).

## Abstract

Rapid and reliable screening of SARS-CoV-2 is fundamental to assess viral spread and limit the pandemic we are facing. In this study we evaluated the reliability and the efficiency of a direct RT-QPCR method (without RNA extraction) using SeeGene Allplex™ 2019-nCoV RT-QPCR and the influence of swab storage media composition on further viral detection.

We show that SeeGene’s assay provides similar efficiency as the RealStar^®^ SARS-CoV-2 RT-PCR kit (Altona Diagnostics), and that RNA extraction is not necessary nor advantageous if samples are stored in UTM or molecular water but is recommended if samples are stored in saline solution and in Hanks medium.

## Introduction

The SARS-CoV-2 pandemic that the world is fighting since January 2020 has already caused devastating mortality (1). As we are facing our biggest public health threat in the modern history, there is a huge number of samples testing to be performed. Most of the current methods described so far are based on viral genes detection by RT-QPCR from patients fluids, including from the respiratory tract (2–4).

To slow the spread of the pandemic, screening is the key (5). This allows to isolate positive patients and stop the transmission. Many medical laboratories now have automated protocol to test the samples. These protocols are based on a Real-time PCR amplification on extracted RNA from fluids or swabs stored in media. However, in many laboratories worldwide, fear of shortage in the chain of supplies is growing. Shortage of RNA extraction kit, PCR detection kits and even swabs storage media will likely arise in many testing laboratories.

This outstanding situation called for alternatives protocols to ensure the continuity of rapid testing in medical laboratories. In addition, the development of new methods could also be used in case of an increase in demand for testing. In this paper, we have compared the concordance rate of positive viral genes detection of extracted RNA from the Abott M2000sp automated plate preparation system using Altona RealStar^®^ SARS-CoV-2 RT-PCR Kit RUO with the SeeGene Allplex™ 2019-nCoV RT-QPCR Assay running on CFX96 BioRad. In addition, we demonstrate that direct testing of viral genes by skipping the RNA extraction step on swabs storage media could be achieved when the samples originate from standard UTM media and from molecular water.

## Methods

Specimens were collected with nasopharyngeal and oropharyngeal swabs, and were stored in different mediums. Mediums used were UTM^®^ (Remel RE12569), homemade Hanks medium, Saline and molecular grade H2O.

Oro-nasopharyngeal swabs were inactivated at 95°C for 5 min and stored at 4°C or −80°C.RNA was extracted from 700 μl of 2 ml of patient’s swab medium using Abbott *m*Sample Preparation Systems DNA kit on m2000sp instrument; the eluate was 90 μl. RT-QPCR plates were automated: prepared on the Abbott m2000sp and routinely detected on m2000rt using the Altona RealStar^®^ SARS-CoV-2 RT-PCR Kit RUO according to the manufacturer’s instruction. 10μl of RNA was used in 20 μl.

SeeGene Allplex™ 2019-nCoV RT-QPCR Assay was manually performed on a CFX96 BioRad machine according to the manufacturer’s instruction. 7 μl of RNA or patient’ swab medium was used in a total of 25 μl and 45 cycles of amplification were performed unless specified otherwise. A patient was considered positive when a positive signal (gene E or S in Altona, genes E, N or RdRP in SeeGene) was detected at any Cts. A patient was considered negative if the internal control was amplified but not the viral genes (N, E, S and RdRP).

## Results and discussion

### Following RNA extraction and RT-QPCR SeeGene vs Altona

To determine the concordance between the standard Altona method and the SeeGene protocol, RNA extracted from 65 SARS-CoV-2-positive and 23 -negative samples were tested using SeeGene kit in three consecutive assays. 23 negative samples were confirmed negative, 64 out of 65 positive samples were confirmed positive (96,92 %). One sample, detected at Ct= 35.6 (gene S only) with Altona’s kit was missed using SeeGene’s kit (Figure 1, Tables 1 and 2). Ct means between the two methods cannot be directly compared as SeeGene uses less RNA, which can have important and this assay was performed following an additional cycle of freeze/thaw. Moreover, both manufacturers target different genes and likely do not use the same primers for the E gene in the PCR which can also results in Ct variation. Although both manufacturers neither use the same amount of RNA in the RT-QPCR reaction nor detect the same viral genes, all compounding factors which could lead to important differences in Ct between the two methods, we still observed a strong correlation between the Ct values obtained by both methods (r=0.9746; *p*<0.0001).

**Figure 1.**
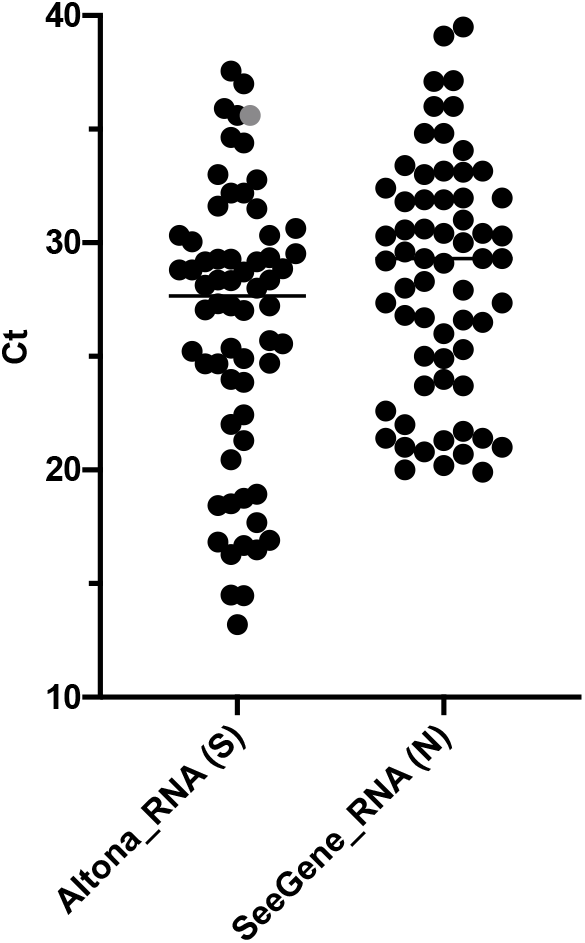
Ranges of Cts detected from RNA extracted from patients Swabs. Each dot represents 1 of 65 reaction, each corresponding to a patient. The grey dot represents the only sample that was missed using SeeGene, detected at Ct= 35.6 with Altona.

**TABLE 1-.**
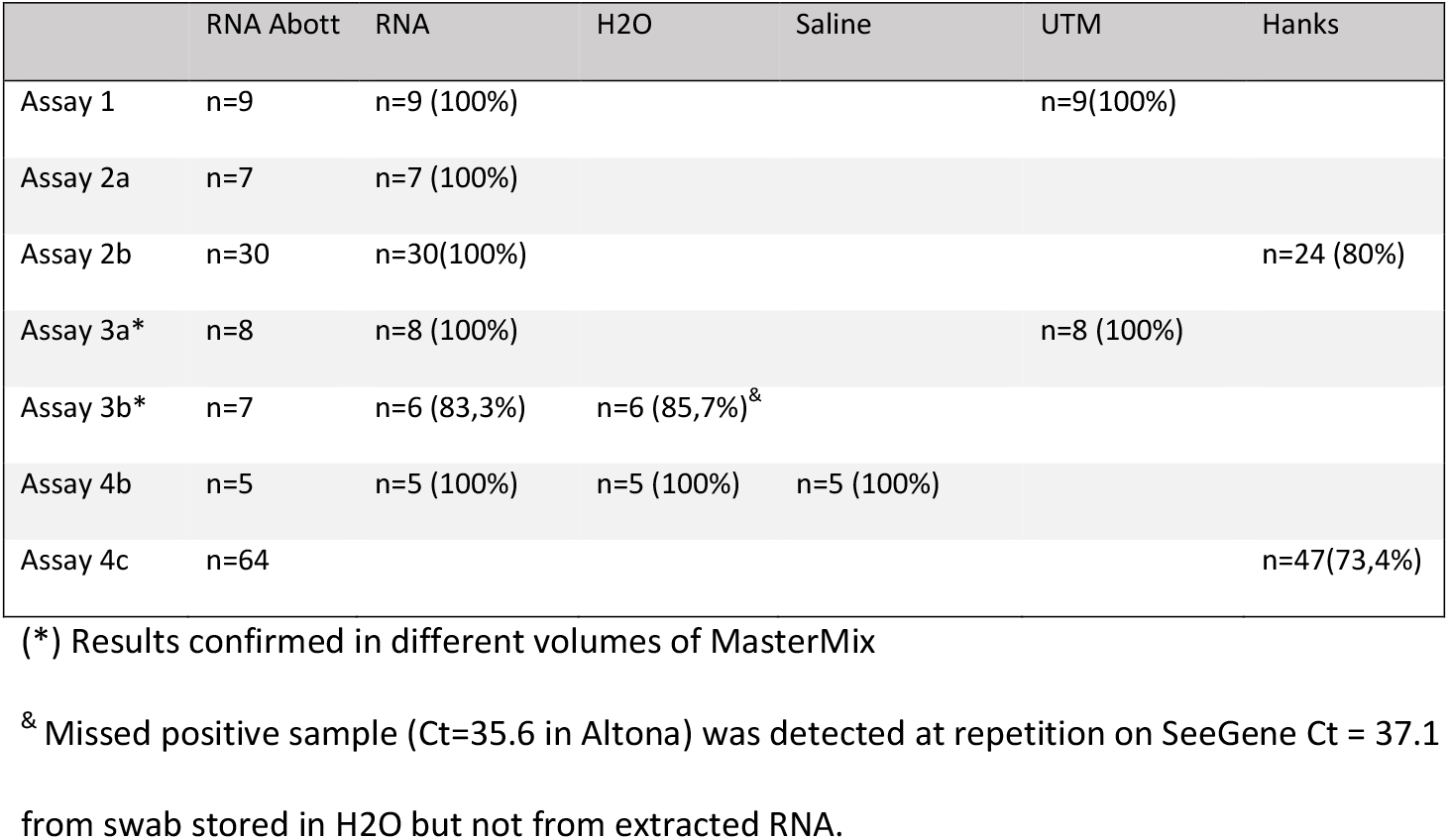
Number of positive samples detected using different methods

**TABLE 2-.** Mean Ct values (targets with best concordance)

|  | RNA Abbott<sup>a</sup> | RNA | H2O | Saline | UTM | Hanks |
| --- | --- | --- | --- | --- | --- | --- |
| Assay 1 | 17,28±6,98<sup>a</sup> | 21,77±8,38 |  |  | 24,22±3,52 |  |
| Assay 2a | 30,65 ±4,32<sup>a</sup> | 31,9 ±4,63 |  |  |  |  |
| Assay 2b | 24,45±4,35<sup>a</sup> | 26,57±5,18 |  |  |  | 31,22 ±6,32 |
| Assay 3a* | 28,75±2,2<sup>a</sup> | 31,15±2,3 |  |  | 30,16±4,6 |  |
| Assay 3b* | 28,02±5,1<sup>a</sup> | 30,73±5,14 | 31,65 ±4,8 |  |  |  |
| Assay 4b | 13,32±0,67<sup>a</sup> |  | 22,6±0,7<sup>b</sup> | 32,98±2,8<sup>b</sup> |  |  |
| Assay 4c | 21,28±5,67<sup>a</sup> |  |  |  |  | 30,68±9,66 |

<sup>a</sup>Abbott Ct values results were added for reference and should not be directly compared to
SeeGene's: RT-QPCRs were performed following an additional freeze/thaw cycle for SeeGene,
less RNA and different primers. <sup>b</sup>These samples originate from RNA eluates diluted hundred
times. (\*) Results confirmed in different volumes of MasterMix.

### RT-QPCR directly from swabs: without RNA extraction

The swab storage media for respiratory virus detection can have important impact on the RT-PCR efficiency, when this media is used directly for PCR reaction, without RNA extraction. To assess the feasibility of direct RT-QPCR using different media we performed the following experiments.

#### Samples stored in UTM

17 samples stored in UTM media, previously assessed as positive by the Altona method with RNA extraction, were used for direct RT-QPCR in two independent experiments. 100% were detected using SeeGene (gene N) and Ct means were equivalent whether or not RNA was extracted before the RT-QPCR (Figure 2; Table 1 and 2). This experiment was also repeated in 12.5 μl of final volume and similar results were obtained.

**Figure 2.**
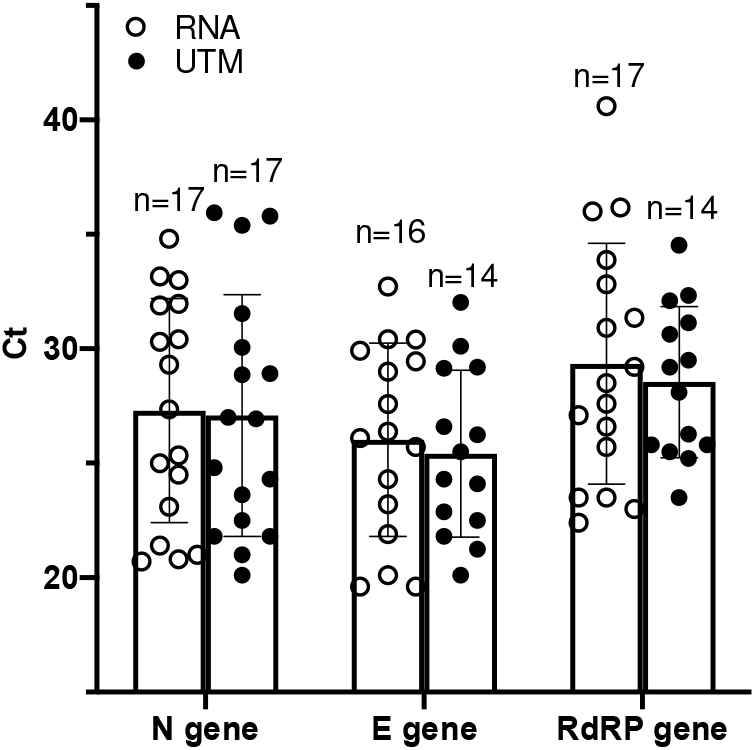
Detection of SARS-CoV-2 by SeeGene RT-QPCR from swabs stored in UTM medium, in the absence of RNA extraction. Each dot represents one reaction corresponding to one patient. Results are plotted as bar graphs with standard deviation. ‘n’ stands for the number of positive RT-QPCR detection.

#### Samples stored in Hanks

Due to a shortage of UTM media, swabs were stored in Hanks medium and analysed for direct RT-QPCR. 94 samples previously assessed as positive by the Altona method (with RNA extraction) stored in Hanks medium were used for direct RT-QPCR in two independent experiments. 21 (2,3%) reactions did not show amplification neither of viral genes nor of the internal control suggesting RT-QPCR inhibition. 75,53% (n=71) were positive following analysis (Tables 1 and 2, Figure 3). The addition of 5 cycles (from 45 cycles to 50 cycles) for 64 of the samples helped with the detection of 5 samples that showed delayed Cts.

**Figure 3.**
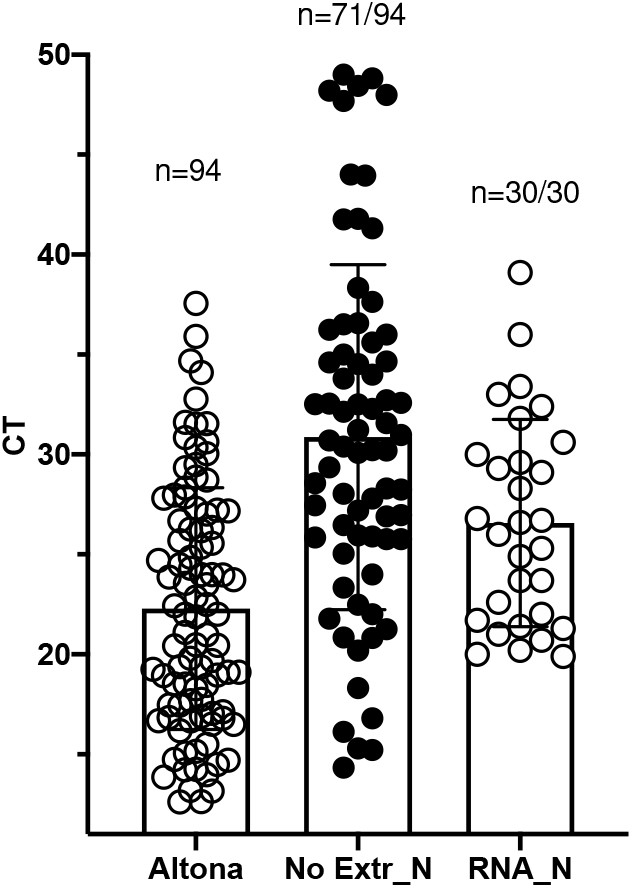
Detection of SARS-CoV-2 from swabs stored in Hanks medium. Results are plotted as bar graphs with standard deviation. Ct values from amplification of extracted RNA from swabs stored in Hanks medium using Altona’s kit (Altona, S gene) and Seegen’s kit (RNA_N; N gene, following an additional freeze/thaw cycle), or directly from Swabs in Hanks medium (No_Extr_N; N gene) using SeeGene’s kit.

Ct values cannot be directly compared to Altona’s, as less volume of sample was added in the mastermix, the assays were performed following an additional cycle of freeze/thaw, and the target genes and primers used by both manufacturers are not the same.

#### Samples stored in water

To determine if molecular grade water could be used as swabs storage media without impacting direct RT-QPCR, three positive swabs were stored in water at 4°C. Eight serial dilutions (1:10) were performed on two of these samples. From the first patient, the undiluted sample and 4 of 7 dilutions showed amplification of SARS-CoV-2 N gene and S gene using RNA with both Altona and SeeGene, respectively. In this same sample comparable amounts of viral N gene was detected without RNA extraction using SeeGene (Figure 4). Cts can be compared between the two methods in this case as there was no additional Freeze/thaw cycle for this experiment. This experiment was confirmed using 12 μl as sample volume in the mastermix. An addition of 5 other patients were efficiently amplified (Figure 5).

**Figure 4.**
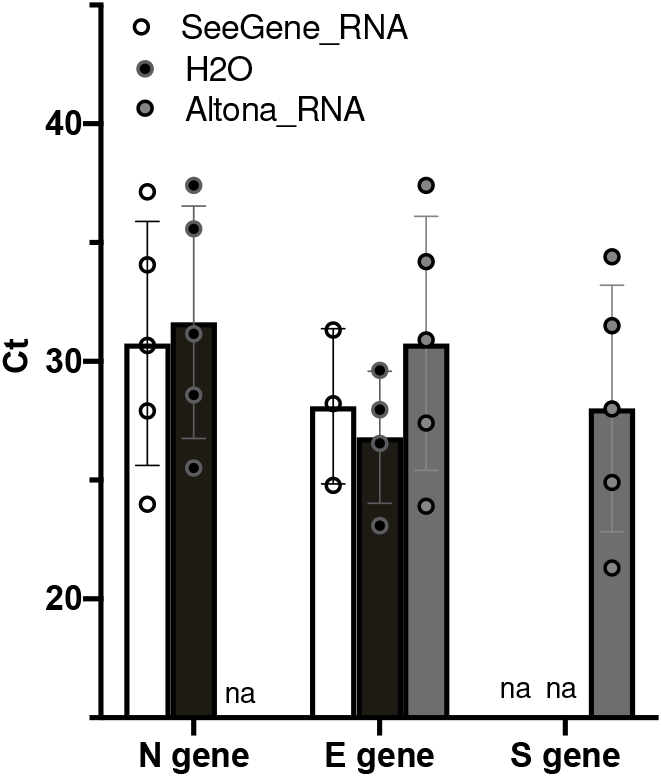
Detection of SARS-CoV-2 by SeeGene RT-QPCR from swabs stored in molecular water. Results are plotted as bar graphs with standard deviation. Serial dilutions were performed in water from a swab stored in water. Ct values from amplification of extracted RNA from swabs stored in water medium using Altona’s kit (Altona_RNA, S and E genes) and Seegen’s kit (SeeGene_RNA; N and E genes), or directly from Swabs in water medium (H20; N and E gene) using SeeGene’s kit. ‘na’ means ‘not available’.

**Figure 5.**
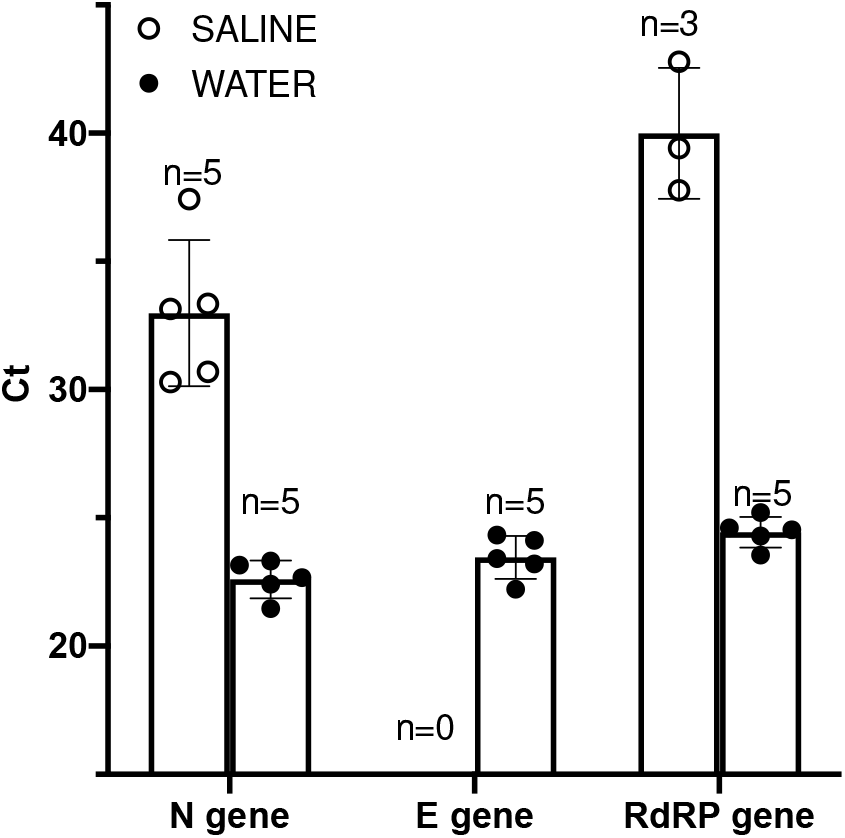
Detection of SARS-CoV-2 by SeeGene RT-QPCR from RNA from swabs diluted in saline water. Results are plotted as bar graphs with standard deviation. One hundredth dilution of 5 samples with high Ct value were performed in molecular water and in saline water. Ct values obtained from RNA eluates diluted in water medium (H2O) or in saline water (Saline) using Seegen’s kit (SeeGene_RNA; RDRP, N and E genes).

**Figure 6.**
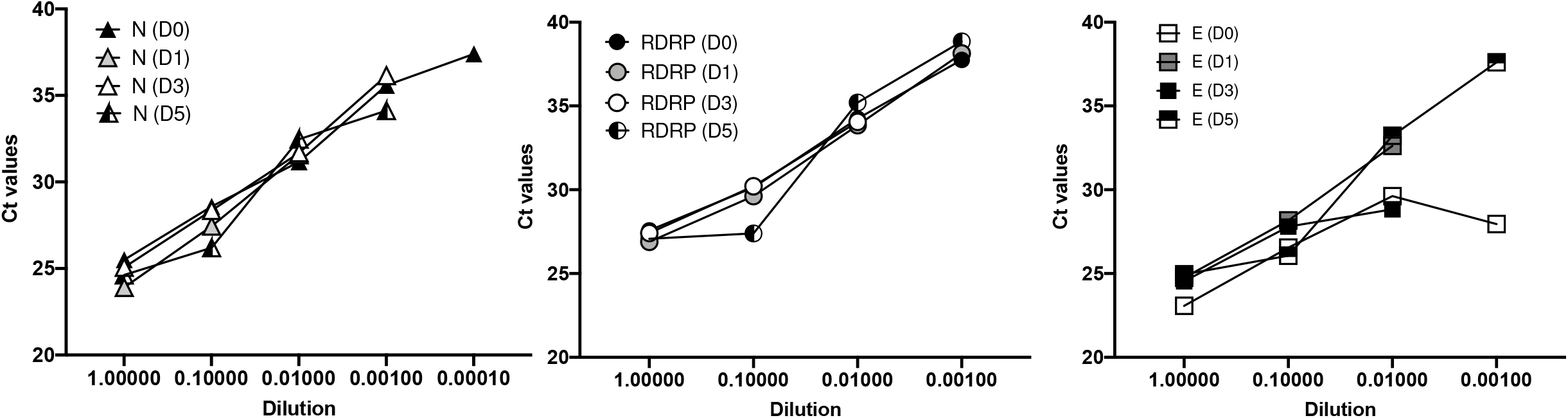
Stability of swabs (not extracted) stored in H2O at 4°C. Serial dilutions were performed in molecular grade water from a swab stored in RNAse inhibitor containing water. Ct values from direct amplification from swab and dilutions was measured using Seegen’s kit (SeeGene_RNA; RdRP, N and E genes) following storage for 0, 1, 3 and 5 days.

In the second sample, SARS-CoV-2 could be detected only from the RNA extracted in the undiluted specimen using Altona’s kit at Ct=35,6. The experiment was repeated using SeeGene and SARS-CoV-2 N gene was detected from H2O (stored at 4°C for 24 hours) at Ct=37.1 but not from extracted RNA (Table 1 and 2).

#### Samples stored in saline solution

To determine if saline solution could be used as swabs storage media without impacting direct RT-PCR, five RNA eluate from samples with high Cts (between 12,61 and 14,22) following Altona’s amplification were diluted in water (1:100) and in saline solution (1:100). Dilutions were directly used for RT-QPCR. As previously observed, amplification could be reliably detected in samples stored in water. However, when samples were stored in saline water, a loss of 10 Cts for the N gene was measured following SeeGene RT-QPCR and the E gene could not be amplified despite 50 cycles.

#### Stability of long-term storage of swabs in water at 4°C

Centralization and increased number of screenings may lead to swabs storage at 4°C for several days. To assess viral RNA stability at 4°C, we quantified levels of N, E and RDRP genes directly from a swab following four 1:10 serial dilutions stored for 0, 24, 72 hours and for 5 days at 4°C. RNase inhibitor was added to the patient swab and serial dilutions in molecular grade water were performed (hence the RNAse inhibitor was also diluted). Viral RNA was detected at 0, 24 and 72 hours at all dilutions but 1:10 000. Ct values were highly constant at all timepoints for the three viral genes for undiluted, 1:10 and 1:100 diluted swabs (E gene, data not shown). The addition of RNase inhibitor probably increased RNA stability of the undiluted swab. Nevertheless, RNA remained highly stable even when diluted hundred times. Samples diluted 1000 times were less accurately amplified: RDRP was detected up to 24h.

## Conclusion

Our results suggest that: i) the SeeGene Allplex™ 2019-nCoV RT-QPCR Assay provides similar efficiency as the Altona RealStar^®^ SARS-CoV-2 RT-PCR Kit RUO, ii) RNA extraction is not necessary if samples are stored in UTM or molecular grade water; iii) RNA extraction is recommended if samples are stored in saline water and in Hanks medium. The number of cycles can be increased to 50 to detect low-producers.

